# First Genome-Wide Centromere Map of *Trypanosoma cruzi* Reveals Linear and 3D Compartment Boundaries and Spatial Clustering

**DOI:** 10.1101/2025.09.26.678825

**Authors:** Natália Karla Bellini, Pedro Gabriel Nachtigall, Pedro Leonardo Carvalho de Lima, Herbert Guimarães de Sousa Silva, David da Silva Pires, Claudia Rabuffo, Danielle Bruno de Carvalho, Yasmin Morais Feno, Ana Paula Cabral de Araujo Lima, T. Nicolai Siegel, Julia Pinheiro Chagas da Cunha

## Abstract

**Background:** *Trypanosoma cruzi*, the etiological agent of Chagas disease, possesses a highly repetitive genome that has historically hindered high-quality assembly and structural characterization. Despite significant advances in assembling *T. cruzi* genomes, major gaps remain. Among these, the complete repertoire of centromeric sequences has remained elusive, representing a critical missing piece in our understanding of chromosome structure and inheritance**. Results:** Here, we generated high-coverage Hi-C (genome-wide chromosome conformation capture) data for the widely used *T. cruzi* Dm28c strain improving its genome assembly, reducing the number of scaffolds and producing a more contiguous and accurate genome. To investigate centromere organization, we performed ChIP-seq using the mNeonGreen-myc-tagged kinetochore proteins KKT2 and KKT3, resulting in the identification of 40 KKT-enriched peaks across 29 scaffolds. These peaks were located in regions enriched in retrotransposable elements, particularly L1Tc and VIPER, near strand switch regions, areas of high GC content, and at the boundaries between conserved genes and virulence-factor multigene families. **Conclusion:** Notably, Hi-C analysis revealed that centromeres may act as structural boundaries contributing to genome compartmentalization and frequently engage in 3D spatial clustering, suggesting a role in higher-order nuclear architecture. Overall, our study provides a high-quality reference genome for the Dm28c strain, presents the first genome-wide centromere map in *T. cruzi*, and offers novel insights into centromere-mediated 3D genome organization

## Background

*Trypanosoma cruzi* is a protozoan parasite and the causative agent of Chagas disease (1), a neglected tropical disease that affects millions of people across Latin America. This organism exhibits polycistronic transcription, lacks canonical promoter sequences (2), and has a genome highly enriched in repetitive sequences. In 2005, the genomes of three trypanosomatids (*T. cruzi*, *T. brucei*, and *Leishmania major)* were sequenced via the Sanger method by a large international consortium, marking the beginning of a new era in trypanosomatid genomics. However, *T. cruzi* genome was the most poorly assembled, likely because of the use of a hybrid strain (CL Brener strain) and its unusually high repeat content compared with those of other trypanosomatids (3). These repetitive sequences represent about 50% of the genome and include multigene families (3–5) as well as various transposable elements (TEs), such as degenerate ingi-related retrotransposons (DIRE-like), tyrosine recombinase (YR) elements, and long interspersed nuclear elements (LINEs) (4). Among LINEs, the L1Tc element is an autonomous retroelement that is present in all *T. cruzi* chromosomal bands (6) and codes for a protein containing four domains (7).

In 2009, forty-one *T. cruzi* chromosomes were assembled *in silico* on the basis of synteny maps and sequences from BAC libraries in the CL Brener strain (8). Since then, advances in genome sequencing and assembly technologies have enabled the sequencing of multiple strains (9–12) from different discrete typing units (DTUs) (13,14). More recently, the use of long-read technologies with the aid of proximity ligation-based methods (Chicago or Hi-C), has led to further improvements in genome assemblies (15). However, very few *T. cruzi* genomes have been assembled at the chromosome level (15,16), and even fewer assemblies resolve chromosome haplotypes (13).

Despite improvements in the genome assemblies of *T. cruzi*, major gaps remain. Among them, the full repertoire of centromeric sequences remains elusive in *T. cruzi*, representing a critical missing piece in our understanding of chromosome structure and inheritance. Only two putative centromeric regions have been described in a very fragmented CL Brener genome assembly (DTU - VI) (17,18). These two putative centromeric regions were based on Topo-II activity and telomere-associated chromosome fragmentation and characterized by the presence of degenerate retroelements, such as vestigial interposed retroelements (VIPERs) and short interspersed repetitive elements (SIREs), as well as transposable elements, high GC content, and localization within divergent (dSSR) or convergent (cSSR) strand switch regions (17,18). In *T. brucei,* centromeres are associated with AT-rich repetitive elements and degenerate retroelements *ingi* and DIRE sequences (19–21), whereas in *Leishmania*, centromeres are essentially nonrepetitive and rich in LmSIDER (large degenerate Ingi-related elements) degenerated retrotransposons (22). Despite differences in centromeric sequences among trypanosomatids, their kinetochore machinery is conserved and composed of unconventional kinetochore proteins, known as KKT (kinetoplastid kinetochore) proteins (22,23). Orthologs of these proteins are also present in *T. cruzi*, but their functional characterization remains largely unexplored.

We recently revealed non-random 3D genomic interactions in *T. cruzi*, including contacts between spliced leader (SL) loci and ribosomal DNA (rDNA) loci, as well as between tRNA genes, indicating a spatial organization that likely optimizes transcription by different RNA polymerase classes (24). In this study, we sought to identify centromeric regions in *T. cruzi* and assess their role in 3D genome architecture, thereby providing new insights into how genome architecture influences gene expression. To this end, we focused on the *T. cruzi* Dm28c strain, a well-established cellular model for *in vitro* differentiation of noninfective forms into infective forms (25), and one for which extensive omics datasets, including RNA-seq, Ribo-seq, tRNA-seq, and proteomics (26–30), are available, enabling comprehensive data integration. Despite its relevance, the most recent genome assembly of the Dm28c strain (DTU I) remains highly fragmented, consisting of 636 scaffolds (9), which limits high-resolution analyses.

Thus, we first improved the genome assembly of the *T. cruzi* Dm28c strain by generating Hi-C data. We then performed ChIP-seq for the kinetochore proteins KKT2 and KKT3 to comprehensively identify, for the first time, the centromeric regions of *T. cruzi*. We show that these centromeres are frequently positioned at borders between the core and multigene family compartments, where they likely act as structural boundaries that contribute to genome compartmentalization. Moreover, we showed that centromeres engage in 3D interactions and cluster within the nucleus, suggesting a key role in higher-order genome organization, potentially influencing gene expression and diversification of surface-virulence factors.

## Results

### Improving the genome assembly of the *T. cruzi* Dm28c strain via Hi-C

The current version of the Dm28c genome available (PRJNA433042, hereafter named Dm28c version 1 - Dm28c.v1), was previously assembled into 636 scaffolds ranging from 0.91 kb to 1.56 Mb, with an N50 of 317.6 kb and a total haploid genome of 53.2 Mb (9). To improve this assembly, we generated Hi-C data from epimastigote forms (**Figure S1**), which provided information about the physical proximity of genomic regions, allowing us to scaffold contigs and enhance the contiguity of the genome assembly. This strategy was previously successfully used in trypanosomes to improve the genome assembly of the *T. brucei* (31), *T. cruzi* - Brazil A4 and TCC strains (15). By integrating the previous genome assembly (SRP134013) with the Hi-C dataset (**Table S1**), we were able to scaffold the initial 636 contigs into 99 scaffolds, and increase the scaffold N50 from 317.6 kb to 866.9 kb, of (**Table S1-2, Fig 1A and Fig S2A)**. The scaffolded assembly (hereafter named Dm28c.v2) revealed an increase in contiguity, with 2.1 Mb for the largest scaffold, in comparison with the 1.6 Mb in the previous assembled version.

**Figure 1.**
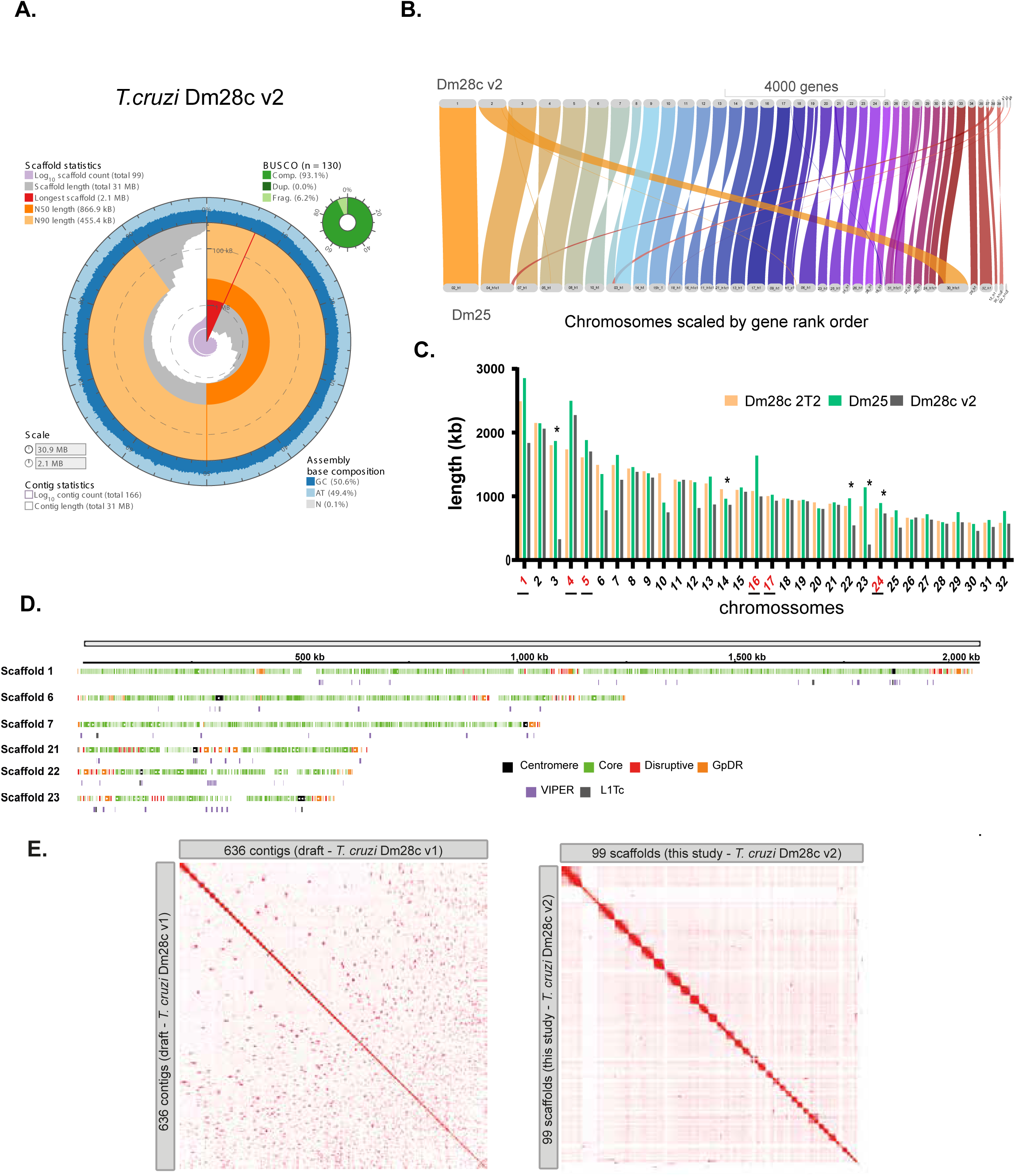
Genome assembly of the *T. cruzi* Dm28c strain via Hi-C reads improves genome contiguity. **A.** Snail plots with genome assembly statistics. **B.** Genome alignment of Dm28c v2 versus Dm25. Scaffolds smaller than 1 kb were not included. **C.** Comparison of chromosome and scaffold lengths (in kb) among Dm28c T2T (32), Dm25 – haplotype H1 (13), and Dm28c v2. Asterisks indicate chromosomes where no centromere was identified. The underlined red numbers highlight chromosomes (based on Dm28c T2T) that are represented by two or more scaffolds in the current assembly. **D.** Linear representation of six scaffolds highlighting the presence of genes from disruptive (TS, mucin and MASP) and GpDR (Gp63, DGF-1 and RHS) compartments at the scaffold ends, likely representing sub and telomeric regions. Centromeres are marked by a black rectangle. **E.** Genome-wide contact maps of Dm28c v1 (636 contigs) and Dm28c v2 (99 contigs), showing fewer trans interactions in the latter as a result of assembly improvements.

Most of the smallest contigs from Dm28c.v1 were composed entirely of tandem repeated DNA sequences (e.g., SL-DNA arrays) with many duplicated contigs, which were purged by our pipeline, resulting in an assembly genome size of 30.9 Mb. Therefore, BUSCO analysis revealed a reduction in the number of duplicates (7.7% to 0%) without major changes in completeness (93.9% to 93.1%) (**Figs. 1A and S2A)**. The quality of both assemblies was also inferred on the basis of the mapping rates of Illumina genomic DNA (gDNA) short reads. Similar mapping percentages (95.5% and 94.3% for v1 and v2, respectively) were obtained (**Table S3**). The completeness of both assemblies was similar, with a striking improvement in the contiguity of v2. In addition, we analyzed the differences in single copy genes, multigene families (MGFs), 18S rDNA loci and other genomic features between assemblies and revealed no significant differences (**Table S3**). Most chromosomes in Dm28c remained diploid in both assemblies (**Figure S2B**), except for scaffold 27, which displayed tetrasomy. This scaffold corresponds to the CL Brener chromosome 31, supporting the notion that its supernumerary status may represent a conserved aneuploidy feature among *T. cruzi* strains.

As expected, comparisons of both Dm28c genome assemblies revealed high identity between versions (**Figure S2C**), except for scaffold 2 which likely consisted of many repetitive sequences from duplicated contigs that were purged in our assembly. Syntenic analysis of their protein-coding genes revealed that many contigs were joined in the improved version, and nine scaffolds showed one-to-one correspondence (scaffolds 8, 24, 34, 36, 37, 38, 40, 41 and 50) (**Figure S2D) (Table S4)**.

As the new assembly proved to be more contiguous, we compared it with the genome assemblies of other *T. cruzi* strains. Notably, our improved genome assembly displays numerous one-to-one syntenic blocks compared with those of the *T. cruzi* Brazil A4 and Dm25 strains (**Figure S2C-D, Figure 1B, Table S5**), indicating high assembly quality. The Brazil A4 genome was assembled via long reads and Hi-C (15), whereas the Dm25 strain presents a chromosome-level phased assembly-based on HiFi reads, composed of 32 pseudochromosomes for haplotype 1 (h1) (13). Impressively, scaffolds 1 to 35, 37-38, 41 and 48 from our assembly correspond to chromosomes 1 to 32 of the Dm25 strain. Notably, scaffolds 8, 9 and 41 form chromosome 3, and scaffolds 27 and 28 form chromosome 31 of Dm25c-h1 (**Table S5)**. Since the Dm25 assembly was generated using long-read sequencing and produced similar results, this further confirms the contiguity and accuracy of our assembly.

While this manuscript was being prepared, a new genome assembly of Dm28c via HiFi reads (referred to here as Dm28c T2T) was deposited on bioRxiv (32). However, the corresponding genome sequence and annotation files were not available, which limited a more extensive and detailed comparison with our assembly. Nevertheless, on the basis of the available synteny data for Dm25 and the reported chromosome lengths, we performed a comparative analysis between our assembly, Dm25, and Dm28c T2T (**Figure 1C and Table S5**). Strikingly, among the 32 chromosomes identified in both assemblies, 28 chromosomes were each represented by a single scaffold in our assembly. Only six chromosomes were composed of two or three scaffolds from our assembly version (**Figure 1C-red numbers**). Additionally, the chromosome lengths of our assembly correspond to approximately 92% (median) of those in Dm28c T2T and 85% (median) of those in Dm25 (**Figure 1C**). Since the Dm28c T2T and Dm25 assemblies were generated using long-read sequencing and yielded consistent chromosomal structures, these findings further support the contiguity and accuracy of our genome assembly on the basis of Hi-C reads.

Gene annotation revealed 11,233 protein coding sequences (CDSs) and 1,051 nonprotein coding genes (**Table S6**). Among the total protein-coding genes, 86.9% were housekeeping genes (core compartment), 8.7% were disruptive genes (trans-sialidases, MASPs and mucins genes) and 4.4% were GpDR genes (Gp63, DGF-1 and RHS genes). As proposed before (32–34), we also noted enrichment of disruptive and GpDR genes at chromosome ends (**Figure 1D**). The noncoding genes represented 1,051 sequences classified as tRNAs, rRNAs, SL-RNAs, snRNA_Us and snoRNAs genes. Overall, we detected a decrease in the number of genes from the previous genome assembly, which likely reflects the purge of duplicated contigs. Finally, telomeric sequences from 17 bp to 3,205 bp were identified in 15 scaffolds. Scaffold 21 contained telomeres at both the 5′ and 3′ ends. Nevertheless, the size of our scaffolds is in line with those proposed for Dm25c and Dm28c T2T (32) (**Figure 1D and Table S7**).

By joining contigs into larger scaffolds, the resulting Hi-C heatmap for previous and improved Dm28c assemblies, indicated a decrease in trans-interactions (intense red rectangles outside the principal diagonal) indicating a correction onto missassembled contigs of the previous assembled version (**Figure 1E**). Compared to the previous version, the Hi-C heatmap from the current assembly displays a more uniform colour gradient, reflecting improved contig ordering and fewer misjoins. This enhanced contiguity allows for a more accurate definition of 3D genome structure, which will be explored below.

### Mapping centromeres in the *T. cruzi* genome

Centromeric DNA has been reported only in the CL Brener strain and restricted to two chromosomes (1 and 3). However, these findings were obtained from a highly fragmented genome assembly, via a telomere-associated chromosome fragmentation strategy combined with topoisomerase-II immunoprecipitation (17,18). To comprehensively determine the centromeric regions in *T. cruzi*, we determined KKT-enriched regions, which are known to discriminate centromeric regions in *T. brucei* and *Leishmania* (22,23,35). Among the 25 KKTs identified in *T. brucei*, we chose KKT2 and KKT3 because they were previously suggested to be associated with *T. brucei* centromeric DNA (23). Thus, we obtained C-terminal mNeonGreen-*myc* (mNG-myc)-tagged cell lines for two *T. cruzi* KKTs (2 and 3) by CRISPR-Cas9. PCR of the genomic DNA confirmed the endogenous tagging of the target genes (**Fig S3A**). Fluorescence microscopy of living epimastigotes revealed intense fluorescence in punctate distributed in the nuclei of either KKT2-mNG-myc (**Fig S3B**) or KKT3-mNG-myc (**Fig S3C**), whereas no fluorescence in the nuclei was observed in the parental line (**Fig S3D**). We then performed ChIP-seq assays using both tagged and untagged parasites (T7Cas9 lineage - control samples). Over 80% of the reads were mapped to the new assembled genome (**Table S8**).

Visual inspection of KKT ChIP-seq coverage revealed a limited number of prominent peaks across the genome, typically one per scaffold (**Figure 2A-C**). To increase stringency, we defined centromeres as regions enriched for KKT2 and/or KKT3 (ChIP/input) exclusively in tagged lineages, excluding any enrichment observed in untagged controls (e.g., ChIP/input of the T7Cas9 lineage). This approach identified 40 significant peaks, ranging from 0.64 kb to 34.87 kb in size (median = 10.2 kb) (**Figure 2B**). The median KKT peak size is in line with the centromeric length of *T. brucei* (from 3.5-54 kb) (21). These significant KKT peaks were distributed across 29 scaffolds (**Table S9**). Among them, 18 scaffolds exhibited a single KKT-enriched region, 9 scaffolds contained two peaks, and 1 scaffold presented three peaks each (**Figure 2B**).

**Figure 2:**
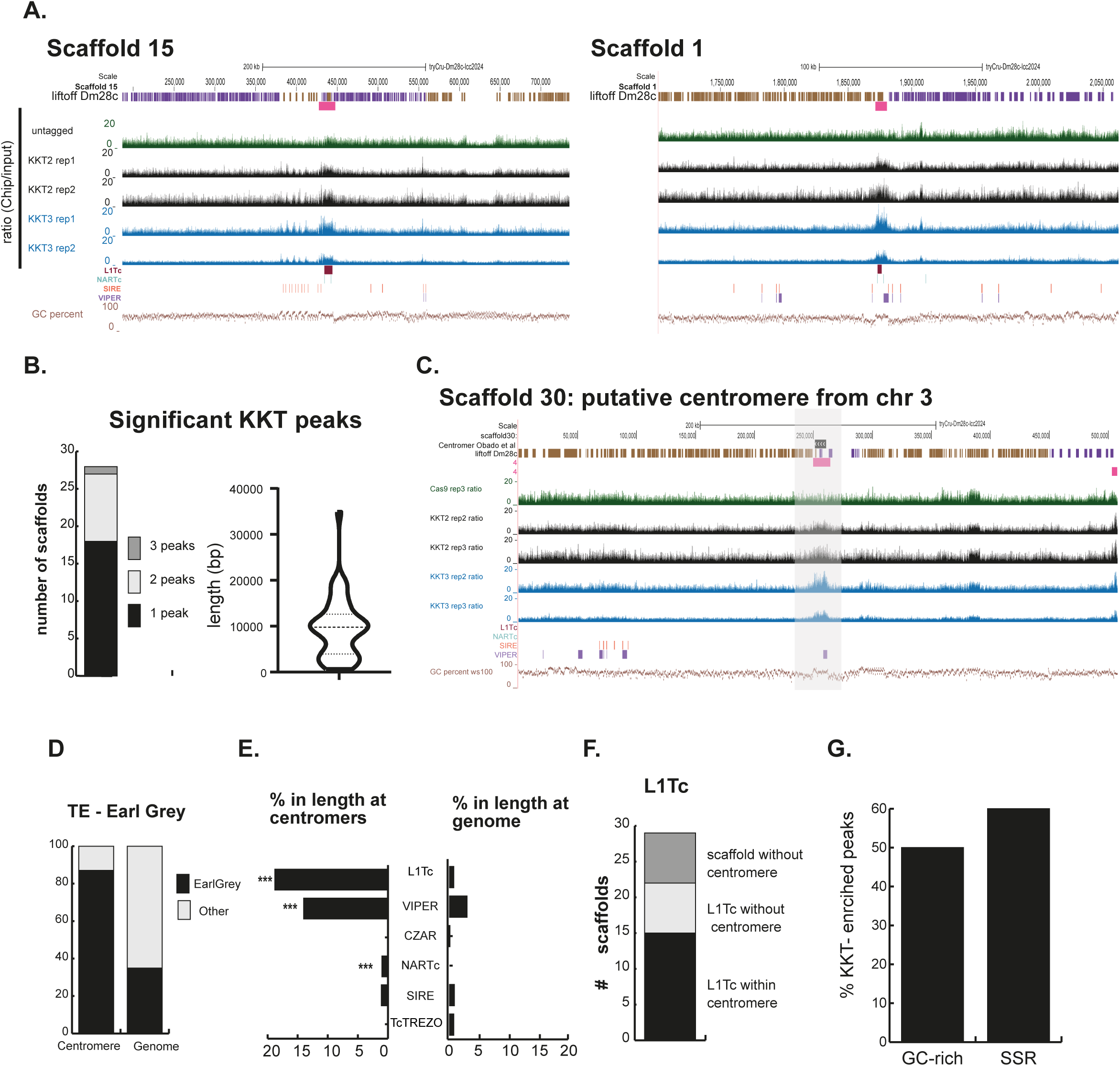
Centromeres of the *T. cruzi* Dm28c genome coincide with L1Tc and/or VIPER retrotransposons. **A.** Integrative Genomics Viewer (IGV) snapshots highlighting the presence of KKT2 and KKT3 peaks as centromere markers. The tracks represent, respectively, the gene annotations (purple for the positive strand and brown for the negative strand), the MACS3 significantly-enriched regions representing the centromeres (pink), the ChIP-seq profiles for the Cas-9 untagged control (green) and KKT2 and KKT3 (black and blue profiles, two replicates each), the retroelements (L1Tc, NARTc, SIRE, VIPER), and the GC profile. **B.** Significant KKT peaks per scaffold (bar graph) and length distribution in bp (violin plot). **C.** KKT2- and KKT3-enriched peaks coincide with genomic regions homologous to the centromeric regions identified in CL Brener (18). Homology analysis of the centromeric sequence from chromosome 3 of the CL Brener strain (earliest assembly) also revealed its presence in Dm28c (black bars). **D.** Proportion of retroelements (by length, based on Earl Grey annotation) in the *T. cruzi* Dm28c genome and centromeres. **E.** Length-based distributions (%) of retroelements in centromeres (left) and in the whole genome (right). **F.** Distribution of scaffolds containing L1Tc elements according to their colocalization with centromeres. **G.** Percentage of KKT-enriched peaks located in GC-rich regions and SSRs.

Among the 32 chromosomes of Dm28c T2T, 27 had putative centromeric regions (**Figure 1B – asterisks and Table S9**). Additionally, we identified a centromeric region on scaffold 29 (**Table S9**), which was not assigned to any Dm25-h1 or Dm28c T2T chromosome, suggesting that it may belong to a distinct haplotype. Among the five chromosomes lacking an identified centromere, chromosomes 3, 22, and 23 were only partially assembled, which likely explains the absence of detected centromeric regions.

To assess whether any of the putative centromeres corresponded to the two centromeric regions previously identified in the highly fragmented CL Brener assembly (chromosomes 1 and 3) (17,18), we performed sequence homology searches against Dm28c v2. Notably, these regions mapped to scaffolds 30 (86% identity and 78.8% coverage) and 29 (92% identity and 62% coverage), respectively, and overlapped with two significant KKT peaks (**Figure 2 C and S4**). Together, these results provide strong evidence that KKT enrichment in *T. cruzi* indeed marks centromeric regions, as observed in other trypanosomatids.

### The retrotransposons L1Tc and VIPER are key centromere marks

We next evaluated whether the 40 KKT-enriched peaks were conserved among themselves and whether they were associated with specific genomic features. To this end, we aligned all the centromeres and observed that both coverage and sequence identity varied considerably, with some centromeres showing little or no conservation (**Figure S5A**).

We further annotated transposable elements (TEs) in the current genome, as they have previously been associated with trypanosomatid centromeres (17–20). A total of 2,694–sequences were classified as TcTREZO, L1Tc, CZAR, NARTc, and SIRE, VIPER, representing 18% of the Dm28c genome (**Figure 2D**). We detected a strong association between KKT enrichment and TEs, especially for the non-LTR retrotransposons L1Tc and VIPER (**Figure 2A, C and D**). On average, 20% of the total length of KKT-enriched regions is composed of L1Tc sequences, followed by 14% corresponding to VIPER elements. Notably, 70% of the KKT-enriched peaks contained L1Tc and/or VIPER elements (**Table S9**).

Although L1Tc is a prominent centromeric marker, not all L1Tc elements are associated with centromeres. For example, centromeres on scaffolds 10, 11, and 16 do not contain L1Tc elements (**Fig S5B and Table S9**), but instead harbor repetitive elements classified as “unknown,” and VIPER/LTR, respectively, according to the Earl Grey annotation tool. In our new assembly, we identified 56 L1Tc sequences (median length: 3,180 bp) distributed across 29 scaffolds (**Table S10**). Among these 29 scaffolds, 22 had an identified centromere, fifteen (15/22 - 68%) colocalized with L1Tc elements, and seven (7/22 - 32%) L1Tc sequences were located outside the region enriched at KKT (**Fig 2F**). We found no major sequence divergence between L1Tc associated with a centromere and L1Tc not associated with a centromere (**Figure S5C)**. Notably, centromeric regions not associated with any TEs presented lower coverage and sequence identity than other centromeric regions did (**Figure S5A-B**).

Tandem repeat searches identified only a few repetitive sequences in centromeres, indicating that *T. cruzi* centromeres are not composed of such elements (**Table S11**). This contrasts with the centromeres of *T. brucei*, which are enriched in AT-rich repeats (20), but resembles the organization observed in *Leishmania* (22).

We found that centromeric regions are significantly depleted in protein-coding genes: only 59 protein-coding genes were identified within centromeric regions, whereas 105 were identified in the control sequences (randomly selected sequences of the same length and within the same scaffold as each centromere). This finding is consistent with our observations that 60% of centromeres are located near or within strand-switch regions (SSRs) (**Fig. 2G**). Finally, 45% of the centromeres were located in regions with elevated GC content (**Fig. 2G** and **Table S9**), and 27.5% were located close to the scaffold ends, likely associated with sub/telomeric regions (**Table S9**) Together, these findings support the notion that L1Tc and VIPER retroelements, strand switch regions, and GC-rich content are conserved features of centromeric regions in *T. cruzi*.

### 3D nuclear architecture in *T. cruzi* mediated by centromere organization

Centromeres have a strong impact on nuclear genome compartmentalization, affecting both intra- and interchromosomal interactions (36). We observed that half of the centromeric regions (20/40) are located at the boundaries between the core and disruptive/GpDR genomic compartments (**Figure 3A–B and Table S9**), raising the question of whether they contribute to the three-dimensional separation of these regions. Correlation matrix analyses revealed that PCA2 of at least four scaffolds (**Figure 3C and S6)** shows a flip coinciding with the position of some centromeres, indicating that the higher-order chromatin structures of their surrounding genomic regions are indeed different. Together, these results suggest that centromeres may act as structural boundaries contributing to the compartmentalization landscape by anchoring or reorienting the chromatin architecture of adjacent genomic regions.

**Figure 3:**
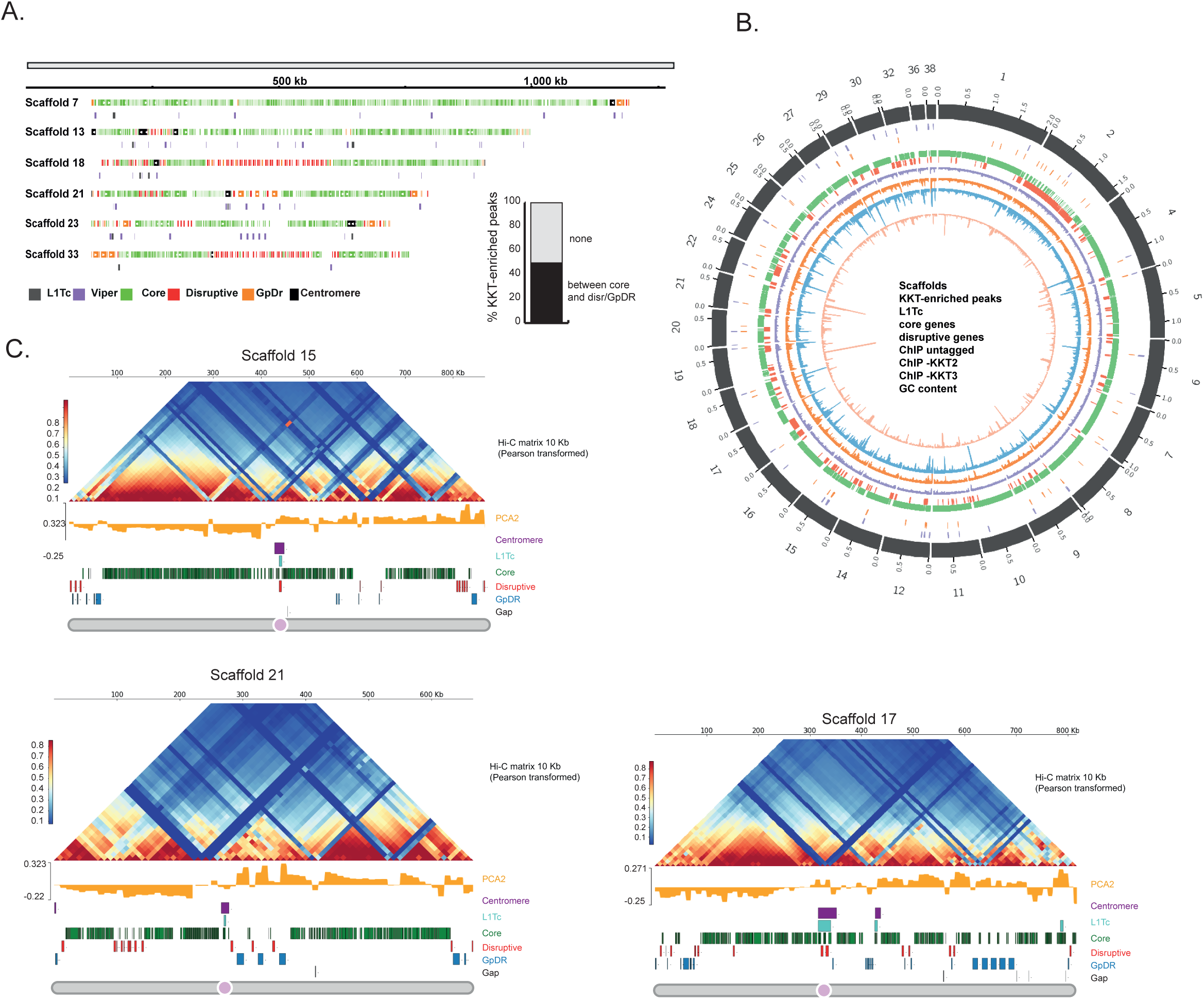
Genomic localization, chromatin context, and 3D positioning of putative centromeres in *T. cruzi*. **A.** Linear representation of six representative scaffolds highlighting the presence of centromeric sequences (black rectangles) between genes from core (green) and disruptive (red), GpDR (orange) compartments. The VIPER (purple) and L1Tc (gray) elements are also depicted. Bar plot showing the percentage of KKT enriched peaks colocalized in between core and disruptive/GpDR regions. **B.** Circus plot of the *T. cruzi* Dm28c v2 karyotype highlighting genomic compartments, centromeres and retroelements mapped along the 29 scaffolds that harbor KKT-enriched peaks. Tracks, from the outer layer to the inner layer, indicate scaffolds (gray), centromeres (purple), L1Tc (orange), core genes (green), disruptive genes (red), ChIP-seq profiles for the control lineage CRISPR-Cas9 (light purple), and the CRISPR-Cas9-tagged kinetochore proteins KKT2 (orange) and KKT3 (blue). The last layer comprises the GC content, in light orange. **C.** Pearson-transformed Hi-C contact matrices at 10 kb resolution highlighting centromere positioning at scaffolds 15, 21 and 17. Principal Component Analysis (PCA) revealed a prominent sign flip in the second eigenvector (yellow track), indicating increased interactions between genomic regions upstream and downstream of the centromeres (purple track). The L1Tc retroelement (turquoise track) shows spatial colocalization with centromeres. Centromeres are frequently located at the interface between chromatin compartments, most often between the core (green track) and disruptive (red track) compartments, or between the core and GpDR (light blue track) compartments.

To further investigate centromere 3D organization, we focused on the 14 scaffolds longer than 1.2 kb that harbor a single centromere. Integration of Hi-C data from epimastigotes revealed that these centromeres interact with each other, highlighted by the distinct contact points in **Figure 4A**. To explore this further, we applied a virtual 4C approach, which enables the assessment of DNA–DNA interactions from a specific locus of interest (viewpoint, VP) across the genome. Consistent with the observations shown in **Figure 4A**, multiple peaks colocalized with centromeres from different scaffolds, indicating 3D clustering of centromeres within the nucleus. Among the centromeres shown in **Figure 4A**, most appeared to interact with each other, except those from scaffold 2 (**Figure 4B**). Importantly, centromere–centromere interactions do not seem to depend on the presence of L1Tc, since centromeres lacking L1Tc (scaffolds 10, 11, and 16) also interact with L1Tc-associated centromeres.

**Figure 4:**
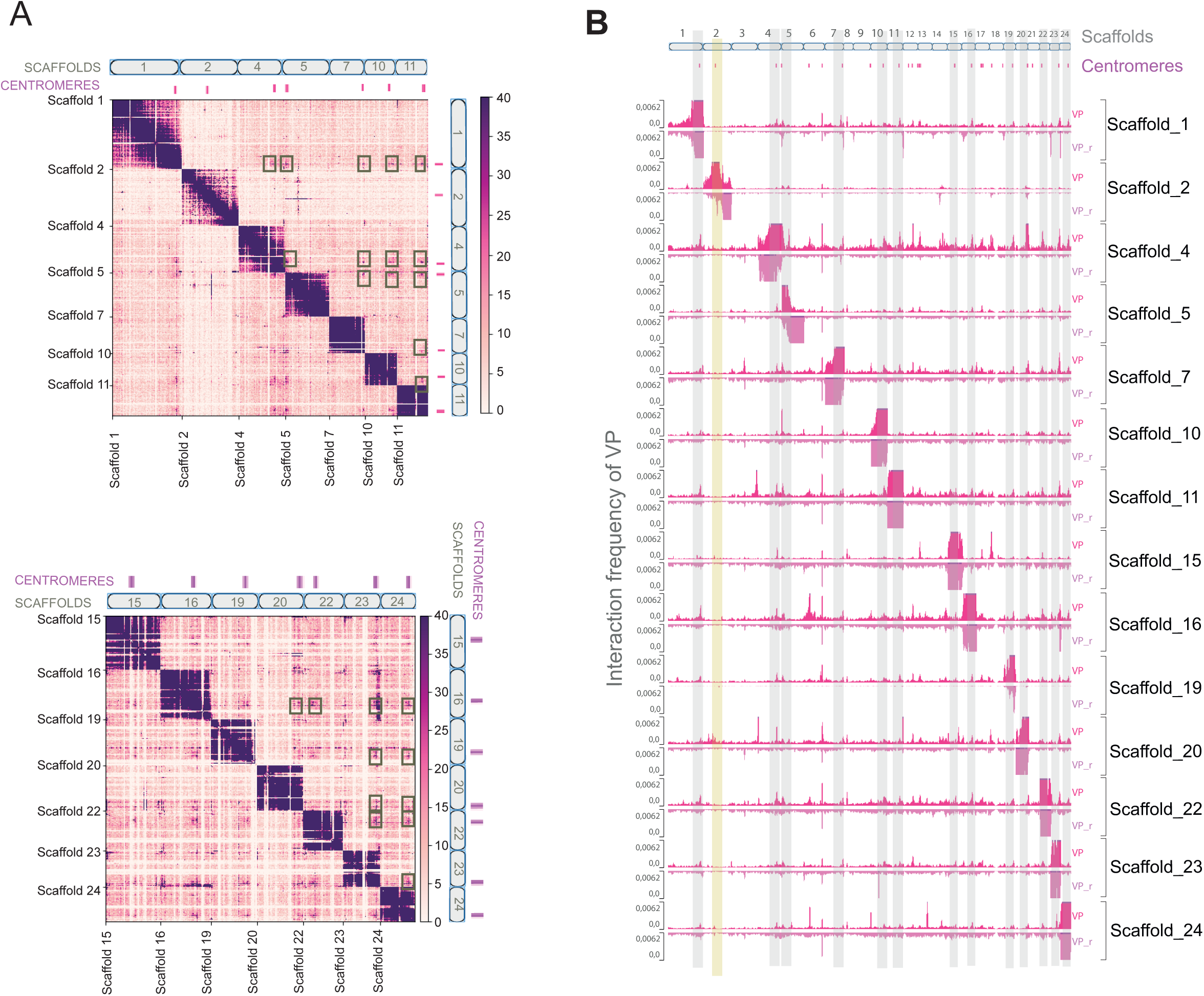
3D clustering of centromeres within *T. cruzi* nuclei. **A.** Hi-C matrix at 15 kb resolution for a subset of scaffolds, each containing a single centromere. The top track (pink rectangles) highlights regions significantly enriched for KKT2/3. The purple square along the main diagonal represents intrascaffold interactions. Light pink signals outside the diagonal indicate trans interactions between different scaffolds. The strong pink dots at these trans interaction zones reflect frequent contacts between centromeres (gray rectangles), suggesting their close spatial proximity in the nucleus. **B.** IGV snapshot of virtual 4C analysis using centromeric coordinates as viewpoints (VPs). The upper plots (pink) display the interaction profiles for each VP across scaffolds 1 to 24. The inverted (light pink) images show control VPs (VP_r), representing interactions of randomly selected regions from the same scaffold as the target centromeric VPs. The vertical shaded bars (gray and yellow) mark the genomic location of each centromere, allowing the assessment of interaction peaks with other centromeres across scaffolds. The colocalization of peaks along these vertical stripes indicates spatial proximity among centromeres, supporting their three-dimensional clustering. The centromere from scaffold 2 (yellow vertical shaded bar) does not interact with other centromeres.

## Discussion

### Hi-C data improve *T. cruzi* Dm28c genome assembly

Obtaining a better genome assembly is crucial to advancing a wide range of studies, including those related to 3D genome organization*. T. cruzi* genome assembly has historically been challenging because of its highly repetitive content (∼50%) (3,4). By integrating Hi-C data with a long-read assembly, we substantially improved the Dm28c assembly, reducing the number of contigs, increasing the N50, and maintaining BUSCO completeness. In addition, the scaffold sizes are in accordance with previous gel band analyses that ranged from 0.57 to 2.58 Mb for *T. cruzi* Dm28c (37) – with the longest scaffold being 2.1 Mb, supporting the reliability of our assembly. The overall genome size (30.9 Mb) was markedly smaller than that in the earlier version, most likely reflecting the removal of duplicated contigs, a common issue in previous assemblies of Dm28c. Nevertheless, the assembly size remains within the range expected compared with that of the closely related Dm25 strain (13).

The synteny observed with Dm25 and Brazil A4 further highlights the accuracy of our assembly, even in chromosomes enriched in disruptive genes, which have traditionally posed greater challenges. The fact that scaffolds corresponding to all 32 Dm28c chromosomes were identified— with lengths approaching the expected sizes—indicates that many of them likely represent complete chromosomes. Moreover, the detection of additional scaffolds potentially representing previously uncharacterized haplotypes points to the value of incorporating Hi-C information to resolve complex regions of the genome. Together, these findings underscore how the integration of chromosome conformation data can not only improve assembly quality but also provide novel insights into the genomic diversity of *T. cruzi*.

### Centromeres in *T. cruzi* are marked by colocalization with L1Tc and VIPER elements

Centromeres are essential genomic loci that anchor the kinetochore, ensuring the faithful segregation of genetic material during cell division. Most eukaryotes are associated with highly repetitive DNA, such as satellite sequences and TEs. For instance, human centromeres are composed of large arrays of alpha satellite DNA interspersed with TEs in pericentromeric regions (38). In trypanosomatids, degenerated retrotransposons have been implicated in centromere composition, with *ingi* and DIRE found at *T. brucei* centromeres and LmSIDER elements in *Leishmania* (22) (**Table 1**).

**Table 1.** Comparison of the main centromeric features of trypanosomatids.

In *T. cruzi,* 70% of centromeres colocalized with L1Tc and/or VIPER elements, suggesting that TEs may contribute to centromere identity. L1Tc, an autonomous non-LTR retrotransposon, has a widespread distribution across nearly all chromosomal bands in the DTUs I, II, V, and VI isolates (37). In the previous Dm28c assembly, 54 L1Tc sequences were mapped across 35 contigs (9). In our new assembly, we identified 56 sequences distributed across 29 scaffolds. Among them, 50% colocalized enriched KKT. In the same sense, 30% of centromeres do not contain L1Tc and/or VIPER elements, suggesting that although these elements are promising centromeric markers in *T. cruzi*, they are unlikely to be the sole sequence associated with centromere identity or function.

Moreover, the presence of retroelements near or within centromeres suggests that recombination events may be facilitated, likely contributing to the diversification and expansion of adjacent sequences. In this context, we found that roughly half of the centromeres are positioned at the boundaries separating regions enriched in conserved genes from regions enriched in surface multigene families, including virulence factors such as trans-sialidases, mucins, and MASPs, which exhibit extensive expansion and diversification in *T. cruzi* (3,4). This strategic positioning of centromeres may have played a role in promoting the diversification of these surface gene families and, therefore, parasite virulence.

In most organisms, centromeres are epigenetically defined by the deposition of CENP-A (a histone H3 variant). The absence of CENP-A in trypanosomatids (39) raises intriguing questions about the signals responsible for KKT recruitment. This task may be particularly challenging, as the centromeric sequences of these organisms appear to vary significantly (**Table 1**). Moreover, the occurrence of more than one KKT-enriched region on the same chromosome suggests that centromere positioning may be flexible, potentially differing among individual parasites within a population.

### 3D organization of centromeres in *T. cruzi* nuclei: putative structural boundary function and spatial clustering

Centromeres have a strong impact on the topological organization of the 3D genome in many organisms, influencing both intra- and interchromosomal interactions. Within individual chromosomes, centromeres can act as barriers, limiting contacts between chromosome arms—an effect that is particularly evident in organisms with Rabl-like nuclear organization, such as *S. pombe* and *A. thaliana*. In addition, centromere clustering contributes to the formation of nuclear subcompartments, which may influence gene expression (40). Here, we found that 50% of the centromeres are positioned between the core and disruptive/GpDR genes, likely separating these groups in three-dimensional space. In addition, in some cases, we observed increased 3D interactions between genomic regions upstream and downstream of centromeres, indicating that centromeres may act as structural boundaries that reorganize local chromatin contacts. These findings support the view that centromeres may contribute to the compartmentalization landscape by anchoring or reorienting higher-order chromatin structures, likely separating regions with different functions and regulations. This is particularly relevant given that core and disruptive/GpDR compartments display distinct 3D genome organization, with the former showing fewer loops (24), longer CFDs (30), more open chromatin (41), and a distinct nucleosome landscape (42). The differential 3D organization of these genomic regions, potentially explained by the presence of centromeres, may in turn contribute to the higher nascent transcript levels (43) observed in core regions compared with disruptive regions in epimastigotes. Whether this organization changes in other life forms, thereby influencing the expression of virulence factors in infective stages, remains to be investigated. Since infective forms are nonreplicative, it will also be important to assess how centromeric regions are arranged in these stages and whether the 3D profiles observed in replicative forms (analyzed here) are maintained.

In various organisms, including plants and animals, centromeres cluster within specific nuclear regions, typically near the nucleolus or at the nuclear periphery, depending on the cell type (40,44). Our analysis of *T. cruzi* nuclear architecture revealed spatial interactions among centromeric regions, which may have functional implications. We identified significant intercentromeric contacts involving multiple centromeric loci across distinct genomic scaffolds, as evidenced by the interaction points observed in both Hi-C matrices and Virtual 4C profiles. Most centromeres appear to engage in such interactions; however, centromere from scaffold 2 does not seem to interact with others, a pattern that does not appear to be explained by their DNA sequence composition.

These findings are consistent with recent observations in *T. brucei*, where centromeres were shown to form nuclear hubs and create subcompartments that separate chromosome arms (21). Similarly, in *Leishmania*, immunofluorescence assays targeting KKT proteins revealed 8–12 distinct nuclear foci—despite the presence of 36 chromosomes—suggesting centromere clustering (22). Taken together, these results indicate that centromere hubs may represent a conserved feature of 3D genome organization, even in organisms with unconventional kinetochores (**Table 1** and **Figure 5**). However, the precise mechanisms and molecular players involved remain to be elucidated. Further studies investigating centromere structure and function across different cell cycle phases will be critical to fully understand the biological relevance of this 3D genomic feature.

**Figure 5.**
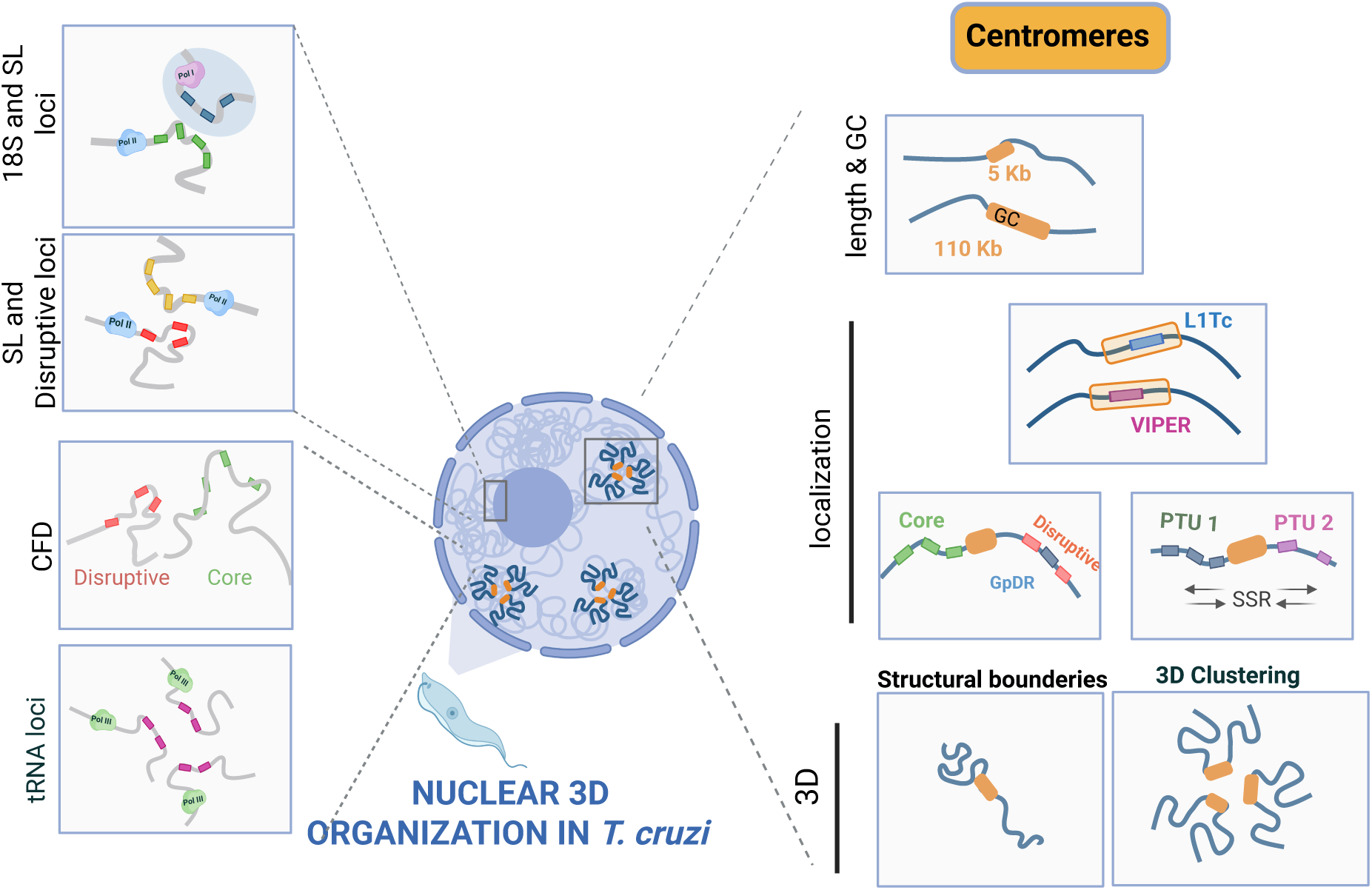
Model of *T. cruzi* genome organization highlighting previous (30,74) and current observations. Highlights on the right summarize the main findings regarding centromeric sequences and their associations with both linear and 3D genome organization. Note that not all centromeric regions exhibit the associations indicated. For a more comprehensive analysis, see Table S9.

## Conclusions

3D genome organization analysis in trypanosomatids provides an excellent system for investigating the impact of this organization on gene expression, since these organisms display little control of transcription initiation (45). Our analysis of 3D genome organization previously showed that, even within this regulatory framework, loci involved in the transcription of essential genes— such as SL RNA, rRNA, tRNAs, and virulence factor genes—are brought into close spatial proximity (Figure 5). Here, we demonstrate that centromeres in *T. cruzi* appear to play multifaceted roles in addition to their classical functions in chromosome segregation. Their preferential localization at boundaries between core and disruptive compartments, together with their clustering in the nuclear space and occurrence in regions of chromatin state transitions (core vs disruptive/virulence factor genes), highlights their potential role in shaping genome compartmentalization. The frequent association of these loci with retroelements such as L1Tc and VIPER further suggests that these loci may act as hubs for genome plasticity, fostering the diversification of multigene families involved in parasite virulence.

## Methods

### Cell culture

*T. cruzi* Dm28c epimastigotes were cultivated at 2 °C in liver infusion tryptose (LIT) medium supplemented with 10% heat inactivated fetal bovine serum and 0.2% glucose, as previously described (46). During exponential growth, 2 × 10□ cells were harvested and processed using the Hi-C wet-lab protocol.

### Hi-C wet-bench protocol

The *in-situ* Hi-C 3.0 experiment was adapted from (47). 200 million epimastigote cells on log-phase were resuspended in 25 mL 1X PBS,crosslinked with 1% formaldehyde solution (50 mM HEPES-KOH pH 7.5, 100 mM NaCl, 1 mM EDTA, 0.5 mM EGTA, 11% Formaldehyde), and incubated for 10 minutes at room temperature with gentle agitation. Fixation was quenched with 750 mM Tris pH 7.5 for 5 min at room temperature, followed by 15 min on ice. Cells were washed four times with 1X PBS, pelleting between washes by centrifugation at 4000 rpm, 4° C for 20 min. For the second cross-linking, cells were resuspended in 6ml 1X PBS, and treated with 3 mM disuccinimidyl glutarate - DSG (50 mg DSG in 511 μL DMSO) for 40 min at room temperature with gentle agitation. The reaction was quenched with 750 mM Tris pH 7.5, 5 min at room temperature, and 15 min on ice. Cells were washed twice with 1X PBS, centrifuged at 4000 rpm, 4 °C, 20 min, and resuspended in 1ml of 1X PBS. To avoid aggregates and cell loss, 0.05% BSA was added, followed by centrifugation at 4000 rpm, 4° C, for 20 min. The supernatant was completely removed and pellet was frozen in liquid nitrogen and stored at −80° C. Samples were thawed on ice and permeabilized with 1 ml of 10 mM Tris-HCl, pH 8.0; 100 mM KCl, 25 mM EDTA, freshly supplemented with 1.4 μM pepstatin A, 4.7 µM leupeptin, 1 mM PMSF, 1 mM TLCK for 5 min at room temperature. Samples were centrifuged at 1800 g, 5 min, 4 °C, and resuspended in 1X NEBuffer 3.1. To increase chromatin accessibility, 1% SDS was added, incubated for 10 min at 65 °C, followed by quenching with 10% Triton X-100 for 15 min, at room temperature. Chromatin was digested with 100 U MboI (NEB, R0147M) and 100 U DdeI (NEB, R0175L), overnight at 37 °C, 400 rpm shaking. Reaction was stopped by incubation at 65°C for 20 min, followed by ice incubation. DNA ends were biotinylated in a fill-in reaction composed of 214 μM biotin-14-dATP (LifeTechnologies, 19524-016), 0.2 mM dCTP, 0.2 mM dGTP, 0.2 mM dTTP, 1X NEBuffer 3.1, 50 U Klenow DNA polymerase I (NEB, M0210), for 4 hours, 23 °C. Proximity ligation with 50 U T4 DNA ligase (Invitrogen, 15224025), in 1X ligation buffer, 1% Triton X-100, 0.1% BSA, for 4 hours, 16 °C in a Thermomixer with interval shaking (900 rpm, 10 s ON, 5 min OFF). Crosslink reversal was performed adding 0.8 mg/mL Proteinase K (Invitrogen, PB107AA) overnight at 65°C Thermal Mixer. DNA was precipitated with ethanol and dissolved in 257 µL of TLE buffer (10 mM Tris-Cl, pH 8.0, 0.1 mM EDTA). Prior to sonication, 0.1% SDS was added, and sample was transferred to microTUBEs Snap-Cap (PN 520045), 130 µL each tube, placed in a Covaris tube holder (#500114), and inserted in a Covaris S220 ultrasonicator (175 peak incident power, 10% duty cycle, 200 cycles per burst, 240 s treatment time). Samples were transferred to an Eppendorf tube, and 0.55X Agencourt AMPure XP beads (Beckman Coulter, A63881) was added to perform the size selection, removing fragments longer than 400 bp. Using 1.3X Agencourt AMPure XP beads fragments smaller than 100 bp were removed by discarding the supernatant, and the target fragments of 200-400 bp remained attached to the beads. The DNA was washed twice in 80% cold ethanol and resuspended in 55 μL 1X TLE buffer. The DNA end repair and biotin removal from unligated ends was performed in a reaction containing the 55 µl eluted DNA, 1X Ligation buffer (NEB, containing ATP), 357 μM dNTP mix, 0.36 U/μl T4 PNK (NEB, M0201), 0.1 U/ T4 DNA polymerase I (NEB, M0203), 0.036 U/ NEB T4 DNA polymerase I, Large (Klenow) fragment (NEB, M0210), 30 min at 20°C, 20 min at 75°C. EDTA (final concentration 10 mM) was added to inactivate the enzymes, and 500 µg Dynabeads™ MyOne™ Streptavidin was used to capture junctions containing biotinylated dATPs incorporated within the chimeric fragments. Beads were washed twice in 400 μl 1X Tween Washing Buffer (TWB: 2.5 mM Tris-HCl pH 7.0, 2.5 mM Tris-HCl pH 8.0, 0.5 mM EDTA, 1 M NaCl, 0.05% Tween 20), resuspended in 400 μl 2X Binding Buffer (5 mM Tris-HCl pH 7.0, 5 mM Tris-HCl pH 8.0, 1 mM EDTA, 2 M NaCl), 330 μl TLE, and 70 μl of the end-repair reaction containing the ligation junction DNA fragments, incubated 15 minutes at RT (400 rpm rotation). Next, 100 μl and 41 μl of 1X TLE buffer were added to wash and resuspend the beads, respectively. 9 μl of A-tailing reaction was added, samples were incubated 30 min at 37° C, followed by 20 min incubation at 65° C. Beads were washed in 450 μl and 400 μl 1X Quick Ligase Reaction and used for Hi-C library preparation.

### Hi-C library preparation and sequencing

Illumina TruSeq adapters were ligated by adding 46.5 μl 1X Quick Ligase Reaction Buffer (NEB, M220), 2.5 μl of DNA Quick Ligase (NEB, M220), and 0.5 μl annealed Illumina TruSeq adapters (50 μM stock), incubated for 1 hour at RT. The beads were washed multiple in 1X NEBuffer 2.1 and 18 μl of DNA samples were used to library amplification with 9 μl of TruSeq PCR primer cocktail (25 μM); 135 μl of 2x KAPA HiFi HotStart Ready Mix (Kapa Biosystems) and 108 μl of water, amplified during 6 PCR cycles following manufacture’s cycling condition recommendations. The product was purified using 1.5X volume of Agencourt AMPure XP beads (Beckman Coulter, A63881), followed by two washes in 70% ethanol, brief air-dry incubation and resuspended in 25 μl TLE buffer. For quality checking 2 nM of the Hi-C libraries were loaded onto a High Sensitivity D5000 ScreenTape®, analyzed in an Agilent 2200 TapeStation system. Based on TapeStation quantification, the sequencing pool was prepared aiming 190 million reads for each sample, and 200X read sequencing depth. The 60 paired-end sequencing was carried out in a NextSeq 2000 Illumina platform as a third-party service at the Gene Center Munich, LMU.

### Scaffolding the Dm28c genome with Hi-C reads

Hi-C data generated in this study and the previously published genome assembly of the *T. cruzi* Dm28c strain (GCA_003177105.1; (9)) were used. The scaffolding process involved removal of duplicates, scaffolding with Hi-C reads, and manual curation to correct misassemblies. Duplicated contigs were removed from the assembled genome using PacBio CLR reads (SRR6809376; (9)) with purge_dups (v1.2.5; (48)), applying mapping parameters specific to PacBio CLR reads in Minimap2 (v2.26-r1175;(49)). The purged genome together with the Hi-C data were used as input for automated scaffolding with YaHS (v1.2a.1; (50)). Manual review of the scaffolded genome was subsequently performed using the Juicer pipeline (v1.6; (51)), following the DNA Genome Assembly Cookbook guidelines (https://aidenlab.org/assembly/manual_180322.pdf). The genome assembly statistics were obtained using the assembly-stats tool (v1.0.1; https://github.com/sanger-pathogens/assembly-stats) and the completeness was assessed using BUSCO (v5.6.1; (52)) with the Euglenozoa gene set (odb10; total of 130 genes).

### Comparative analysis

Comparisons between the newly scaffolded genome and the previous Dm28c v1 assembly, Brazil A4 genome (GCA_015033625; (15)), and the Dm25c genome (PRJNA1039287; (13)) were performed using GENESPACE (v1.3.1; (53)). Data from the Dm28c T2T assembly (Supplementary Material deposited at bioRxiv; (32)) were also incorporated to estimate chromosome length and assess synteny with the Dm25c genome. Genomes were also compared using the dotPlotly tool (https://github.com/tpoorten/dotPlotly). Genomes were mapped using Minimap2 (v2.30;(49)) with default parameters. Dotplots were obatined using the script “pafCoordsDotPlotly.R” from genome alignments.

### Structural genome annotation

To annotate the scaffolded Dm28c.v2 genome, we transferred the gene annotations from the genome assembly of the *T. cruzi* Dm28c.v1 GCA_003177105.1 (9); using liftoff v1.6.3 (54). For non-coding RNA genes we used the sequence of ncDNAs annotated in Dm28c.v1 and we performed BLASTn searches using the “findbestmatch” script (https://github.com/trypchromics/tcruzi-origami), as previously described (24). The gene annotation file was used as input to the “annotatePolycistron” script (https://github.com/alexranieri/annotatePolycistron) to annotate the polycistrons of coding sequences, as well as the extragenomic regions transcription start and termination sites. Transposable elements were identified using the EarlGrey pipeline (v5.0.0; (55)), executed with default parameters. NARTc, SIRE, CZAR, VIPER, and TcTREZO elements annotated in Dm28c.v1 were mapped to the Dm28c.v2 assembly using BLASTn (56). Only hits with 100% coverage and ≥85% sequence identity were retained. Redundant or overlapping hits were merged using bedtools merge (book-ended merge). L1Tc sequences obtained from the Dm28c v1 assembly (9) were aligned against the Dm28c v2 assembly using BLAST, and hits with at least 90% identity and 90% coverage were retained. Hits within 400 bp were merged using bedtools *merge*. Telomeric sequences at scaffold termini were identified using tidk-search v0.2.0 (57)), with TTAGGG as query.

### Quality genome assembly and chromosome copy number

Genomic DNA (gDNA) obtained for epimastigotes (Dm28c) (in biological duplicates) was sequenced on the Illumina platform at the Instituto Butantan. The paired-end short reads (2 x 150) were aligned against the reference genomes TriTrypDB-Dm28c-2018 (v42) and LCC-Dm28c-2024 (versions v1 and v2, respectively) using Bowtie2, with customized parameters optimized for local and sensitive alignment (--local -D 25 -R 4 -N 0 -L 19 -i S,1,0.40 --n-ceil L,0,0.15 --dpad 15 --gbar 4 -X 800). To estimate chromosome copy number (CCN), a pipeline based on (58) that integrates alignment data and coverage-based analysis was created. Coverage and comparative analyses were performed using bamCoverage and bamCompare from deepTools (59), with RPKM normalization across curated set of loci derived from single-copy genes (4,775 genes) obtained from (9) defined genomic bins of 500 pb, effective genome size of 30.872.613 pb. The mean scaffold coverage over single-copy genes was computed using BEDTools (60) to infer relative copy number by scaffolds. Final CCN values were normalized against reference chromosomes and summarized in a tab-separated values (TSV) file. A custom Python script to visualize the chromosomal CNV (copy number variation) profiles was used (Git-hub).

### Genome editing of *T. cruzi* targeting the KKT2 and KKT3 genes

Epimastigotes of *T. cruzi* Dm28c constitutively expressing T7 RNA Polymerase and SpCas9 were used as a parental line (61) to generate endogenously tagged lines as previously described. The LeishGEdit tool (62) was used to design primers considering the genome of Dm28c (available at Table S12) for the generation of DNA templates for 3’ guide RNAs, and of repair cassettes, by PCR using the pPOTc plasmid containing the mNeonGreen (mNG) gene followed by 3 repeats of the myc sequence, and either the *BSD* gene (resistance to blasticidin) or the *PAC* gene (resistance to puromycin), and the Q5 High Fidelity DNA Polymerase (NEB, USA). PCRs to generate repair cassettes were conducted using primers DFW (either KKT2_DFW or KKT3_DFW) and REV_TAG (either KKT2_REV_TAG or KKT3_REV_TAG), and conducted at 98°C (5 min), followed by 35 cycles of denaturation at 98°C (30 sec), annealing at 55°C (30 sec) and extension at 72°C (2:30 min), followed by 5 min extension at 72°C. PCRs to generate RNA guide templates were conducted using primers 3GUIDE (either KKT2_3GUIDE or KKT3_3GUIDE) and G000, at 98°C (5 min), followed by 30 cycles of denaturation at 98°C (30 sec), annealing at 60°C (30 sec) and extension at 72°C (30 sec), followed by 5 min extension at 72°C. PCR products for repair cassette and for the 3‘guide template were pooled and purified using the PCR purification kit (QIAGEN, Germany). Epimastigotes of the T7/Cas9 parental line were collected at mid-log phase, washed once in transfection buffer (90 mM Na_2_HPO_4_, 5 mM KCl, 0,15 mM CaCl_2_, 50 mM Hepes, pH 7.4), resuspended in 200 μL of the same buffer, and incubated for 2 min at room temperature with the DNA mix. The mixture was transferred to electroporation cuvettes of 0.2 cm gap (VWR Scientific, Pennsylvania, USA), placed in an 2D Amaxa equipment (Lonza, Basel, Switzerland) and pulsed once using the U-033 program. Parasites were transferred to LIT supplemented with 10% FCS and cultivated for 24 h before addition of blasticidin at 25 μg/mL for the selection of KKT2_mNG_3xmyc or puromycin 50 μg/mL for the selection of KKT3_mNG_3xmyc. Cultures were selected for 25 days, and the genomic DNA of the selected populations were purified using the DNeasy extraction kit (QIAgen, Germany). Integration was checked by PCR using 100 ng of gDNA as a template, and primers KKT2_FW_checkTAG and KKT2_REV_TAG or KKT3_FW_check_TAG and KKT3_REV_TAG using GoTaq (Promega, Madison, WI) according to manufacturer’s instructions.

### Fluorescence microscopy

Epimastigotes were collected by centrifugation at 700 x g for 10 min and resuspended in RPMI supplemented with Hoechst 33342 (R&D systems, USA) diluted 1:2000, and incubated at 28°C for 20 min. An aliquot of the solution was collected and mounted on a glass slide, covered with a round coverslip previously coated with poli-L-lysine (Sigma). Live parasites were subsequently imaged at a Cell Observer Yokogawa Spinning Disk (ZEISS) laser confocal microscope, under 100X magnification oil-objective.

### ChIP-seq assay mediated by the KKT2 and KKT3-myc lineages

ChIP-seq assay was performed following (63) with minor modifications. Briefly, 3×10^8^ *T. cruzi* Dm28c epimastigote myc-KKT2, myc-KKT3 and the untagged lineages expressing the Cas9 (all in three biological replicates), were cross-linked with 1% formaldehyde solution in phosphate buffered saline (PBS), and 1 mM phenylmethylsulfonyl fluoride (PMSF) during 5 minutes at room temperature. The reaction was stopped with 125 mM glycine, and the parasites were then resuspended in LB1 buffer (50 mM HEPES-KOH pH 7.5, 140 mM NaCl, 1 mM EDTA, 10% (vol/vol) glycerol, 0.,5% (vol/vol) NP-40/Igepal CA-630 e 0.,25% (vol/vol) Triton X-100, 1 mM PMSF), homogenized for 5 minutes, and washed in LB2 buffer (10 mM Tris-HCL (pH 8.0), 200 mM NaCl, 1 mM EDTA and 0.,5 mM EGTA). The pellet was then resuspended in 500 µL TELT buffer (50 mM Tris–HCl pH 8.0, 62.,5 mM EDTA, 2.,5 M LiCl, 4% Triton X-100) and transferred to 0.5 mL sonication tubes (PCR-0.,5-L-C-Corning). The sample was sonicated with a Qsonica Q800R3 sonicator (50% amplitude, 15 s ON, 30 s OFF, 30 min, at 4 □). G Dynabeads (Thermo Fisher #10003D) were resuspended in blocking buffer (5 mg/mL BSA in PBS) containing 4 µg of anti-myc antibody (Cell Signaling Technology #9B11) and incubated at 4°C overnight. At this step, 10 µL of the previously sonicated supernatant was retained to be used as input sample, as control. Then, the remaining fraction (ChIP sample) was subsequently incubated with the antibody-bead complex overnight at 4°C under gentle rotation, followed by eight washes with cold RIPA buffer (50 mM HEPES-KOH pH 7.5, 500 mM LiCl, 1 mM EDTA, 1% NP-40, 0.7% Na-deoxycholate, 1mM PMSF), and a final wash with TE buffer (10 mM Tris-HCl pH 8.0 e 1 mM EDTA pH 8.0). The sample was eluted in 50 mM Tris-HCl pH 8.0; 10 mM EDTA pH 8.0, and 1% SDS at 65°C for 30 minutes. Cross-links in the ChIP and input samples were reversed with 300 mM NaCl (65° C, 17 hours) and treated with RNase A (Thermo Fisher #EN0531) and proteinase K (Thermo Fisher #25530049). After DNA purification by Phenol Chloroform method, both DNA samples (ChIP and input samples) were used to construct an Illumina library using TrueSeq adapters and sequenced by Illumina NextSeq 1000 with 150 bp by paired end.

### ChIP-seq data processing and centromere identification

Raw ChIP data were evaluated by fastQC (64), adapters were removed and reads were trimmed (crop 99, headcrop 14, leading 0, trailing 0, maxinfo 85:0.05 and minlen 35) using trimmomatic (65) with parameters set to “crop 99, headcrop 14, leading 0, trailing 0, maxinfo 85:0.05 and minlen 35”. The trimmed reads were mapped against the Dm28c v2 genome assembly using Bowtie2 v.2.4.5 (66) with parameters set to “*local*, -N 0 -L 19 -i S,1,0.40”. To obtain KKT2, KKT3 and control (from untagged Cas9 lineages) peaks, ChIP and input bam files were loaded using the *callpeak* tool from MACS3 v3.0.2 (67) with the following parameters “-g 30872613 -broad --broad-cutoff 0.01 -B -q 0.01 --max-gap 1000”. The intersect of the KKT2 and KKT3 ChIP/input peaks (276 peaks) and KKT2-ChIP/Cas9-ChIP and KKT3-ChIP/Cas9-ChIP peaks (80 peaks) were obtained resulting in 44 significant KKT peaks. Manual curation removed peaks located entirely within scaffold 99 or near assembly gaps, and peaks separated by less than 20 kb were merged, resulting in 40 significant KKT peaks. BigWig files were generated using the BAMCoverage tool from deepTools (59) with default parameters, except for the options “--effectiveGenomeSize 53271887” and “--normalizeUsing RPGC”. The BigWig files from ChIP and input samples were compared using bigWigCompare with the options “--operation ratio” and “--binSize 1”, considering three replicates. ChIP-seq data visualization was performed using the Integrative Genomics Viewer (IGV; http://software.broadinstitute.org/software/igv/) and the UCSC Genome Browser.

### Centromeric sequence analysis

The nucleotide sequences of the two centromeres identified by Obado et al. (2005, 2007) (17,18) at chromosomes 1 and 3 were mapped onto the Dm28c v2 genome assembly using BLAST (blastn 2.16.0+) with options enabled to calculate both percent identity and hit length. The latter was used to compute the coverage of the aligned region, which in turn was employed to rank and subsequently select the best match. The nucleotide sequences corresponding to the 40 significant KKT peaks were ranked by length and aligned using BLAST (Align two or more sequences; web interface version available in July 2025). The longest sequence was used as the initial query against all other sequences as subjects, and this procedure was repeated iteratively until the penultimate sequence was aligned against the last one. The resulting coverage and identity percentage values were then summarized and displayed in a heatmap. Tandem repeats were identified in centromeric regions using Tandem Repeats Finder (v4.09; (68)) with default parameters (2 7 7 80 10 50 500).

### Preprocessing of Hi-C reads and DNA-DNA contact matrix construction

Hi-C reads were evaluated with fastQC tool (64). Adapters and low-quality nucleotides were trimmed and reads lower than 35 nts were removed using Trimmomatic (65). The mHi-C pipeline (69), which is extensively described at (24) and accounts for the inclusion of multimapping reads associated with repetitive regions, was used for mapping filtered Hi-C reads onto *T. cruzi* Dm28c v2 reference genome. The resulting alignments file was used to create the contact matrices (.mcool) and to normalize using the Knight-Ruiz (KR) algorithm (70). Matrices of 5 kb and 15 kb resolution were used for downstream analysis with Virtual4C approach (using 5 kb matrix), as well as for matrix visualization through Juicebox (71) and HicExplorer (72).

### Scanning centromere mediated interactions by Virtual-4C

Virtual 4C analysis was performed with the Hi-C Sunt Dracones (HiCSD) package (https://github.com/foerstner-lab/HiCsuntdracones) as previously described (24). Viewpoints (VPs) were defined as the centromere coordinates midpoint ±5 kb (10 kb total). Control VPs of equal size were randomly selected from the same scaffolds using bedtools (random function) (60). The pipeline outputs were visualized in IGV (https://igv.org/app/). Additionally, Circos v. 0.69-8 (73) was used to plot a circular genome consisting of eight tracks including the ChIP-seq profiles of centromeres and L1Tc repeats.

## Declarations

### Ethics approval and consent to participate

Not applicable.

### Consent for publication

**Not applicable.** This study does not involve human participants or identifiable personal data.

### Availability of data and materials

The raw Hi-C reads are available at the NCBI SRA -PRJNA1321695. The genome assembly has been deposited at DDBJ/ENA/GenBank under the accession JBREBH000000000.

### Authors’ contributions

NKB performed the Hi-C wet- and dry-bench analyses, with important contributions from CR and TNS. PGN developed the DM28c v2 genome assembly, annotation, and comparative analyses. DBC, YMF, and APCAL generated the KKT2 and KKT3 cell lineages. NKB and HGSS carried out the KKT ChIP-seq assays. PLCL, DSP, and JPCC processed and analyzed the ChIP-seq data and retroelement annotations. APCAL, TNS, and JPCC supervised students and secured the funding used to perform the experiments. The manuscript was primarily written by NKB and JPCC, with valuable input from all authors. All authors read and approved the final manuscript.

## Acknowledgments

We thank Anna Barcons-Simon for establishing the Hi-C protocol in Siegel’s lab, Raúl O. Consentino for support with demultiplexing during Hi-C data analysis, and Kellana Santos e Silva for assisting with the preparation of crosslinked extracts from tagged strains and for initial studies on putative centromeric regions and their association with Hi-C data in *T. cruzi* Brazil A4. We also thank Leticia Lopes for providing *T. cruzi* gDNA extracts, and Vinicius Carius De Souza for valuable suggestions. We are grateful to all members of the TrypChrOMICs laboratory for many insightful discussions. We thank Dr. Loyze Lima and the Laboratório Estratégico de Diagnóstico Molecular at the Butantan Institute for ChIP-seq NGS analysis, and the Gene Center Munich at LMU for sequencing the Hi-C libraries. This work utilized resources from the High-Performance Computing System of the Center for Bioinformatics and Computational Biology (NBBC) at the Butantan Institute.

## AI STATEMENT

This manuscript was reviewed with the assistance of ChatGPT for grammar, spelling, and clarity improvements. All scientific content, interpretations, and conclusions remain at the sole responsibility of the authors.

## Supplemental Material

### Supplemental Figure Legends

**Figure S1:** Hi-C wet-lab quality control protocol. **A.** Electrophoretic analysis of DNA products (50 ng per lane) on 5% non-denaturing polyacrylamide gels, stained with SYBR Gold (Thermo Fisher). MM - molecular marker 100 bp DNA Ladder (NEB #B7025); qc1 - cross-linked DNA; qc2-digested chromatin; qc3-ligated DNA, qc4-fragmented DNA after sonication; qc5-size selection fractions comprising target (T), upper (U) and lower (L) fragments. **B.** Library amplification titration (QC6). PCR products obtained after 6, 8, 10, 12, and 14 amplification cycles were electrophoresed on 5% non-denaturing polyacrylamide gels and stained with SYBR Gold (Thermo Fisher). In accordance with the Hi-C 3.0 protocol recommendations (Lafontaine et al., 2021), 6 cycles were selected for the assay. **C.** TapeStation report analysis for Hi-C library electrophoresed on a High Sensitivity D5000 ScreenTape (Agilent). Graph (right side): lower (15 bp) and upper (1000 bp) peaks stand for the High Sensitivity D5000 Sample Buffer reference markers. The middle peak shows the fragment size average (421 bp) of the T.cruzi Hi-C library (A1) seen at the electrophoreses image (left panel).

**Figure S2:** Comparative genome assembly evaluation and synteny analyses of *T. cruzi* Dm28c v2. **A.** Snail plots of genome assembly statistics from *T. cruzi* - Dm28c v1. **B.** Ploidy analysis based on the genome coverage of single-copy genes obtained from Illumina short reads after mapping gDNA reads onto both assemblies versions. **C.** Dot plot analysis comparing genome assemblies of Dm28c (v1 versus v2, left) and Dm28c v2 with Brazil A4 (right). **D.** Synteny analysis of *T. cruzi* assemblies: Dm28c (v1 versus v2) and Dm28c v2 versus Brazil A4. Scaffold synteny analyses were performed using coding DNA sequences (CDSs).

**Figure S3.** Genome editing at KKT2 and KKT3 loci by CRISPR Cas9. **A.** Agarose gel of PCR products confirming genome editing at KKT2 and KKT3 loci. The oligonucleotide sequences are described in Table S12. The cruzipain (czp) PCR products were used as a positive control. **B.** Fluorescence images of live epimastigotes expressing mNG-myc-KKT2 and mNG-myc-KKT3.

**Fig S4.** The enriched peaks KKT2 and KKT3 coincided with genomic regions homologous to the centromeric regions identified in CL Brener (17,18). Homology analysis of the centromeric sequence from chromosome 1 of the CL Brener strain (earliest assembly) also revealed its presence in Dm28c (black bars). ChIP-seq profiles for the Cas-9 untagged control (green) and KKT2 and KKT3 (black and blue profiles, two replicates each), retroelements (L1Tc, NARTc, SIRE, VIPER), and GC profiles.

**Figure S5:** Sequence similarity, retroelement association, and phylogenetic relationships of KKT-enriched centromeric regions in *T. cruzi*. **A.** Heatmap of identity (upper triangle - brown) and coverage (lower triangle - blue) analysis of 40 KKT-enriched peaks. **B.** Summary plots of the BLASTN analysis of L1Tc sequence (dm28c v1 - PRFA000001: 1559757-1564634 - 4877 bp - subject – red bars) and centromeric regions from representative scaffolds 1, 2, 4, 5, 7, 15, 19, 20, 22, 23, and 24 (query – blue bars). For each analysis, the percentages of query coverage (Q Cov) and identity (P Ident) are depicted. Centromeres from scaffolds 10, 11 and 16 do not align with the L1TC sequence and therefore are not represented. **C.** Phylogenetic tree of genomic sequence of 40 KKT-enriched peaks.

**Figure S6:** Pearson-transformed Hi-C contact matrices at 10 kb resolution highlighting centromere positioning at scaffold 25. Principal Component Analysis (PCA) reveals a prominent sign flip in the second eigenvector (yellow track), indicating increased interactions between genomic regions upstream and downstream of the centromeres (purple track). The L1Tc retroelement (turquoise track) shows spatial colocalization with centromeres. Centromeres are frequently located at the interface between chromatin compartments, most often between the core (green track) and disruptive (red track) compartments, or between the core and GpDR (light blue track) compartments.

## Supplemental Tables

Table S1: HiC metrics

Table S2. T. cruzi Dm28c v2 genome assembly statistics compared to the previous Dm28c v1 and available assemblies for DTU I strains.

Table S3. Comparison of gDNA read mapping percentages between T. cruzi Dm28c genome assemblies v1 and v2.

Table S4. Correspondence of scaffolds from Dm28c v1 into v2.

Table S5. Correspondence of scaffolds from Dm28c v2 into Dm28c 2T2 and Dm25.

Table S6. Number of protein-coding and non-coding genes in four genome assemblies from T. cruzi - DTU 1.

Table S7: Telomeric DNA in T.cruzi Dm28c genome, its genomic position and length (in bp).

Table S8: KKT2 and KKT3 ChIP-seq metrics.

Table S9: Centromeric features of T. cruzi.

Table S10: L1Tc coordinates at Dm28c v2 assembly.

Table S11: Tandem repeat finder output of 40-KKT-enriched peaks.

Table S12: Oligonucleotides used to obatin and check edited cell lines.

