## Supplementary figures and images for "First Genome-Wide Centromere Map of *Trypanosoma cruzi* Reveals Linear and 3D Compartment Boundaries and Spatial Clustering"

### Supplemental Figures

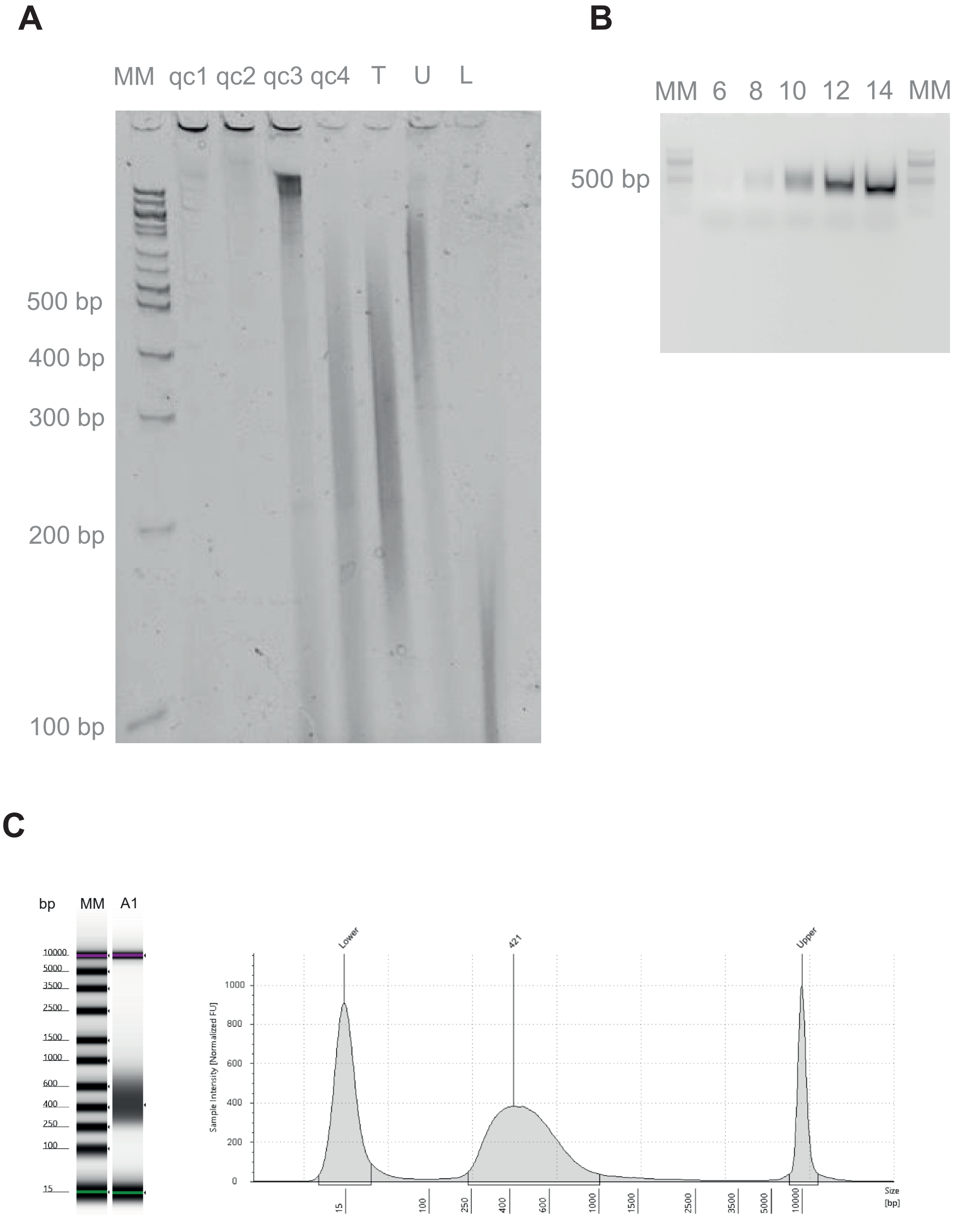

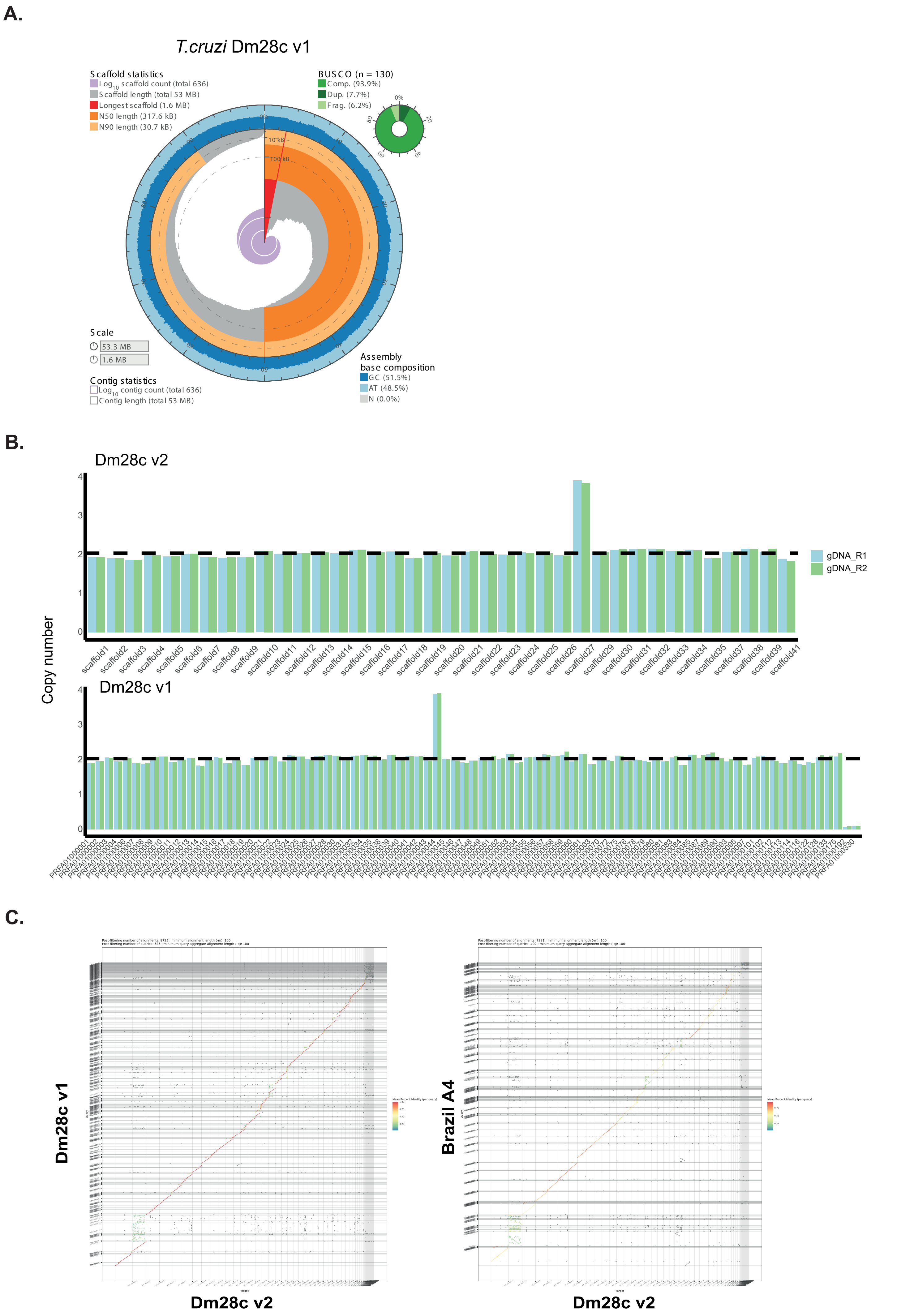

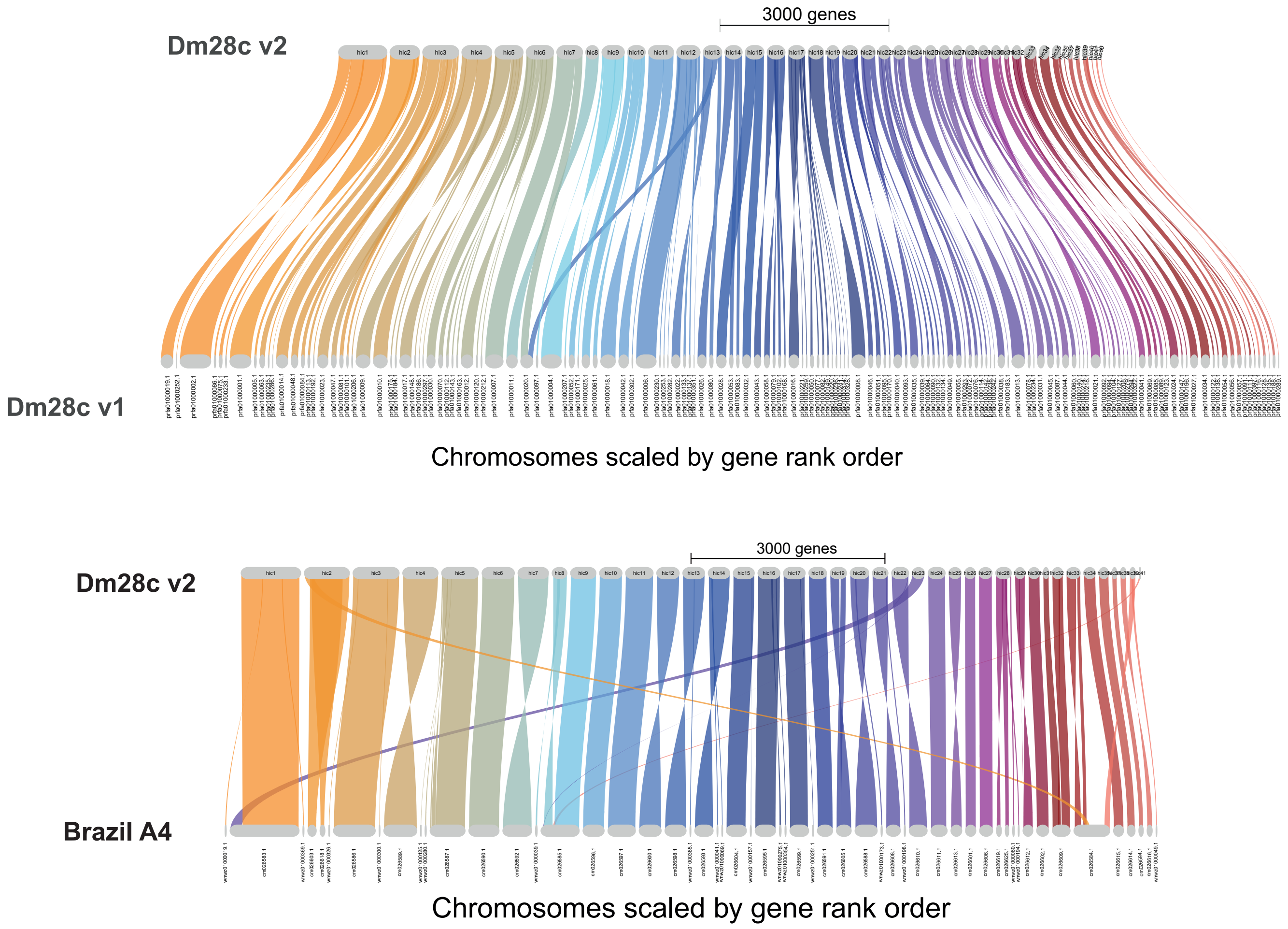

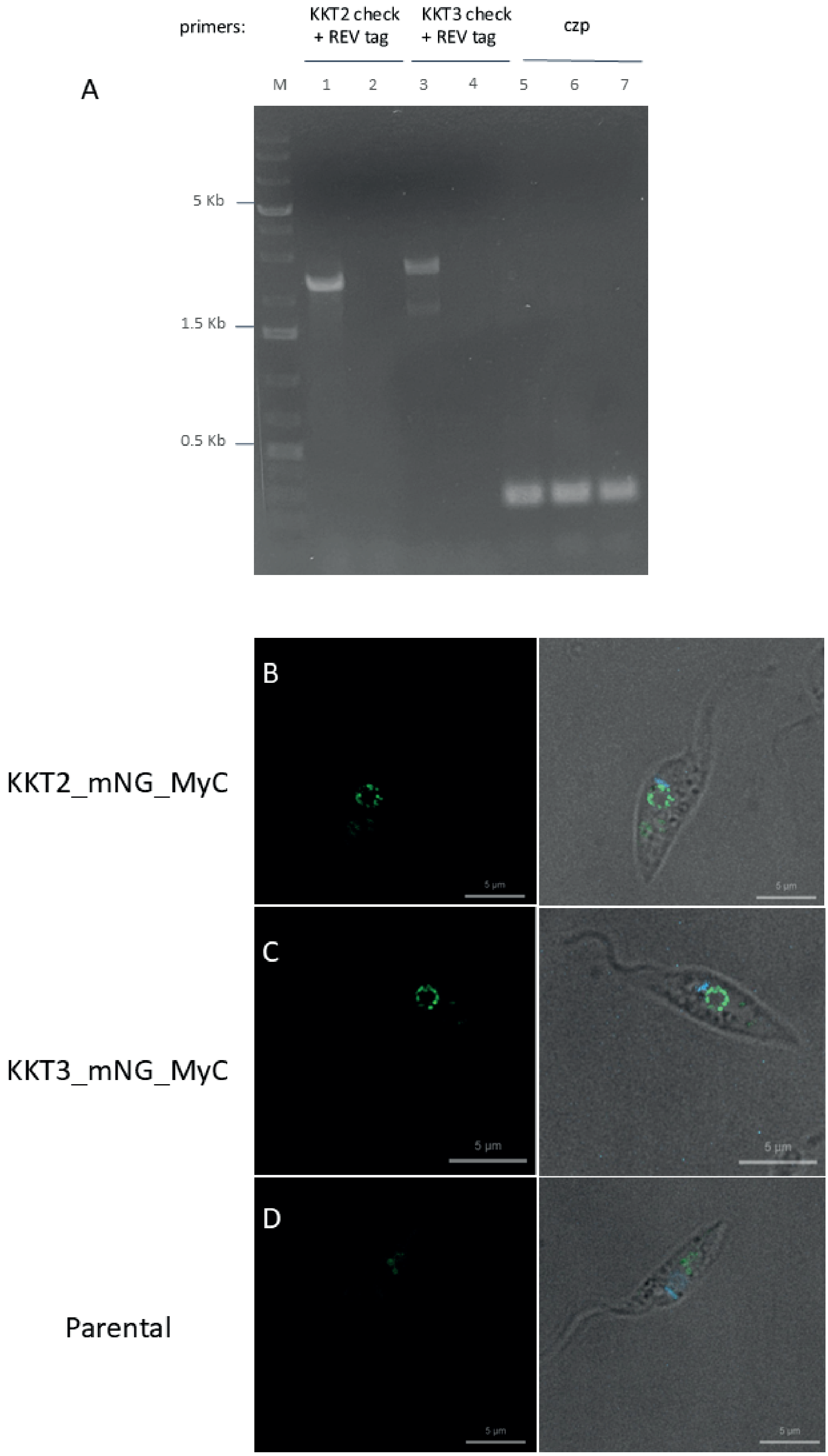

#### Scaffold 29: putative centromere from chr 1

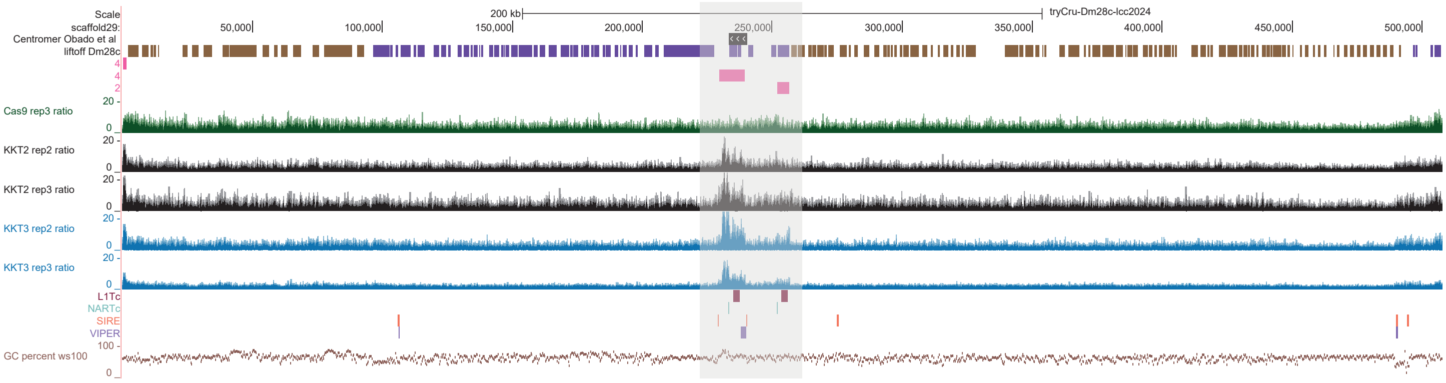


Figure S4

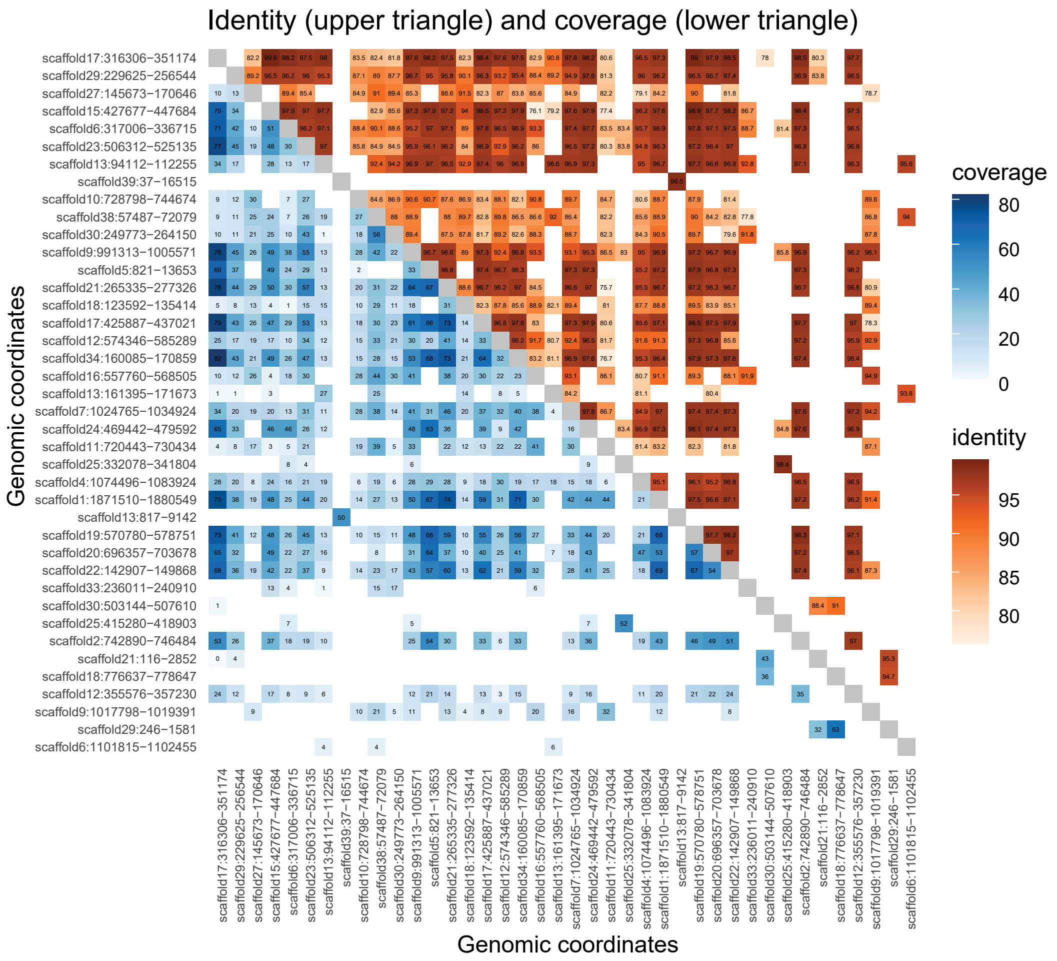

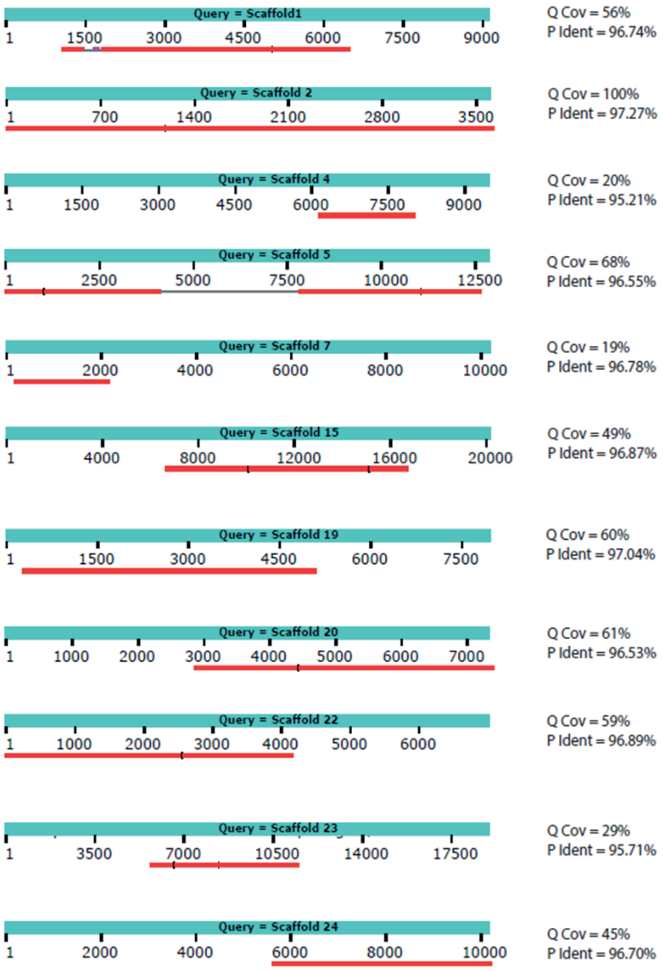

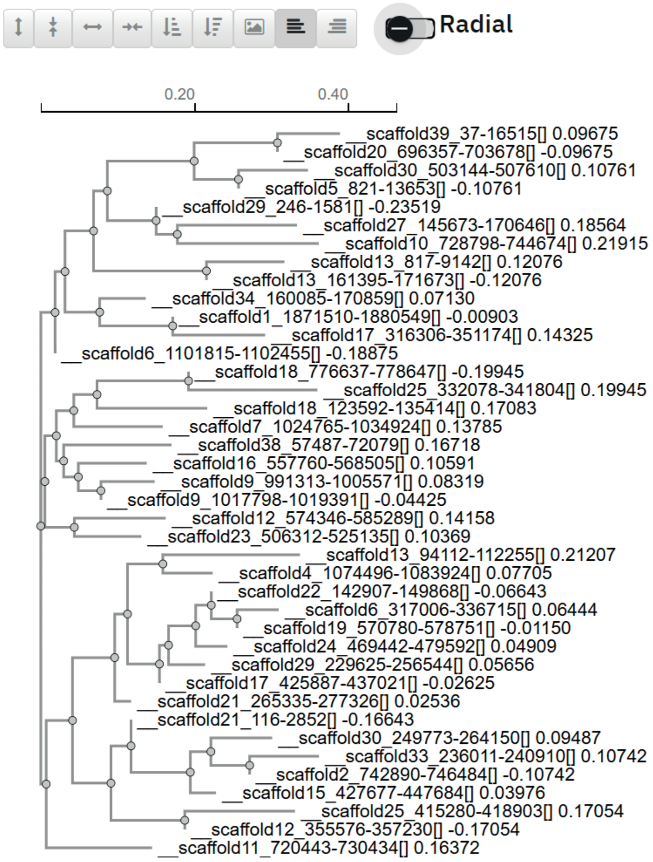

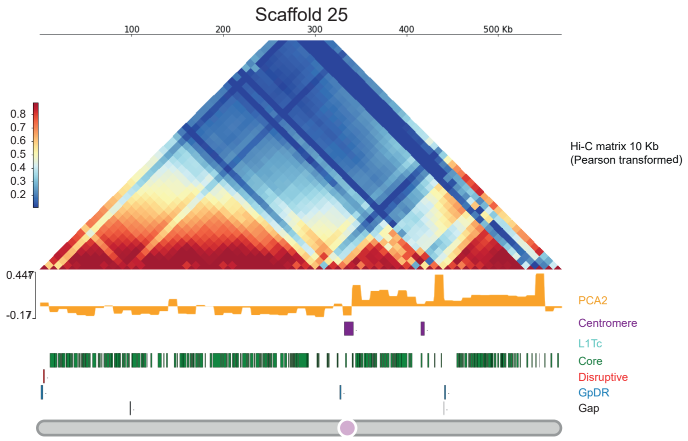
